# Genetically Encoded Boronolectin as a Specific Red Fluorescent UDP-GlcNAc Biosensor

**DOI:** 10.1101/2023.03.01.530644

**Authors:** Jing Zhang, Zefan Li, Yu Pang, Yichong Fan, Hui-wang Ai

## Abstract

There is great interest in developing boronolectins, which are synthetic lectin mimics containing a boronic acid functional group for reversible recognition of diol-containing molecules, such as glycans and ribonucleotides. However, it remains a significant challenge to gain specificity. Here, we present a genetically encoded boronolectin, which is a hybrid protein consisting of a noncanonical amino acid (ncAA) p-boronophenylalanine (pBoF), natural-lectin-derived peptide sequences, and a circularly permuted red fluorescent protein (cpRFP). The genetic encodability permitted a straightforward protein engineering process to derive a red fluorescent biosensor that can specifically bind uridine diphosphate N-acetylglucosamine (UDP-GlcNAc), an important nucleotide sugar involved in metabolic sensing and cell signaling. We further characterized the resultant boronic acid-and peptide-assisted UDP-GlcNAc sensor (bapaUGAc) both in vitro and in live mammalian cells. Because UDP-GlcNAc in the endoplasmic reticulum (ER) and Golgi apparatus plays essential roles in glycosylating biomolecules in the secretory pathway, we genetically expressed bapaUGAc in the ER and Golgi and validated the sensor for its responses to metabolic disruption and pharmacological inhibition. In addition, we combined bapaUGAc with UGAcS, a recently reported green fluorescent UDP-GlcNAc sensor based on an alternative sensing mechanism, to monitor UDP-GlcNAc level changes in the ER and cytosol simultaneously. We expect our work to facilitate the future development of specific boronolectins for carbohydrates. In addition, this newly developed genetically encoded bapaUGAc sensor will be a valuable tool for studying UDP-GlcNAc and glycobiology.

---

Carbohydrates constitute one of the four major classes of biomolecules, and they play a crucial role in many physiological and pathological processes. As such, specific carbohydrate binders are highly sought after for their potential applications in research, diagnostics, therapy, and drug delivery.^1-3^ In recent decades, researchers have focused on developing carbohydrate-binding antibodies, lectins, aptamers, and synthetic lectin mimics.^4-6^ Despite these efforts, the development of specific binders for biologically significant carbohydrates remains a significant challenge in the field.

Boronic acids can undergo reversible esterification reactions with molecules that contain cis-diol functional groups (Fig. 1a).^7^ This property allows boronic acids to recognize and bind to carbohydrates, leading to the development of synthetic lectin mimics known as “boronolectins”. However, these interactions are often promiscuous and lack specificity. Researchers have attempted to improve the specificity of boronic acid-carbohydrate interactions by using multiple coordination sites, steric effects, and molecular geometry, but these approaches have only been successful for a few examples.^6,8-10^ A generalizable approach is still lacking.

**Figure 1.**
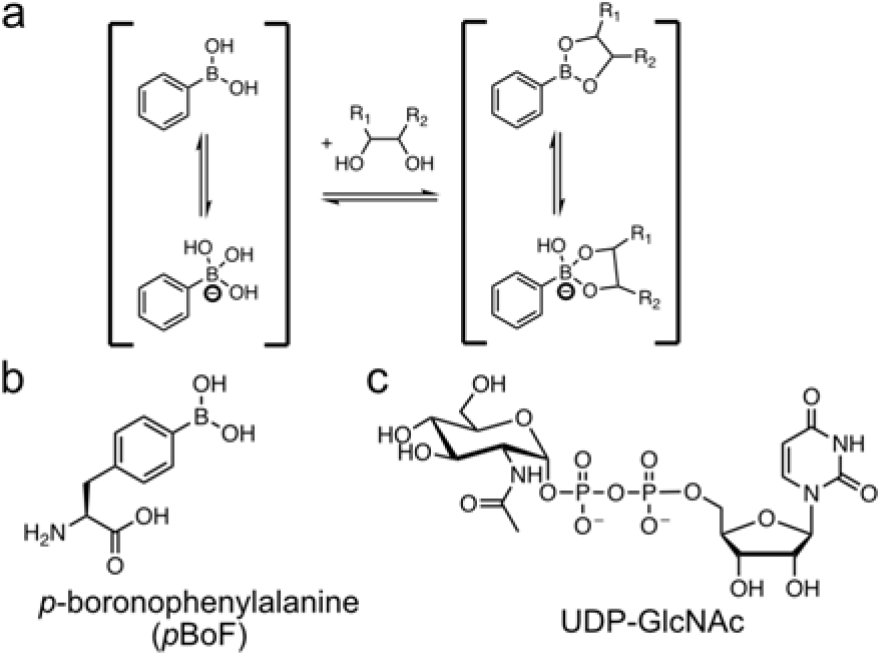
(a) Chemical reaction between phenylboronic acid and diol. (b, c) Chemical structures of the indicated compounds.

Genetic code expansion (GCE) is a technology that allows for the convenient expression of proteins containing noncanonical amino acids (ncAAs) in live cells and organisms.^11,12^ GCE uses engineered tRNA and aminoacyl-tRNA synthetase to introduce ncAAs into proteins in a site-specific manner. Previous studies have enabled the genetic encoding of p-boronophenylalanine (pBoF, Fig. 1b) in E. coli and mammalian cells.^13,14^ In addition, pBoF has been introduced into a phage display library, leading to the identification of pBoF-containing single-chain variable antibody fragments (scFvs) that bind to glucosamine (GlcN).^15^ However, the specificity of these pBoF-containing scFvs to glucosamine, compared to other structurally similar carbohydrates, was not sufficiently addressed in this previous study.^15^

Fluorescent protein (FP)-based biosensors have become popular tools for biological and medical research because they can provide real-time and specific detection of a wide range of biomolecules and bioactivities.^16^ Introducing ncAAs into FPs is an attractive strategy for developing new FP-based biosensors.^17,18^ Previously, we and others incorporated pBoF into FPs and developed reaction-based biosensors for hydrogen peroxide and peroxynitrite.^14,19-21^ Despite the progress, pBoF-modified FPs have not yet been used to sense glycans.

Based on the knowledge and results described above, we hypothesize that pBoF-modified FPs may be engineered into specific biosensors for carbohydrates. Because GCE provides a practical method for the genetic encoding of pBoF-modified FPs, specificity may be achieved via genetic protein engineering. To test our hypothesis, we selected uridine diphosphate N-acetylglucosamine (UDP-GlcNAc, Fig. 1c), an important cell metabolite, as our first target.

UDP-GlcNAc is considered an integrator of nutritional and metabolic signals.^22^ In mammalian cells, UDP-GlcNAc can be synthesized de novo from glucose, glutamine, acetyl-coenzyme A (Ac-CoA), adenosine triphosphate (ATP), and uridine triphosphate (UTP) via the hexosamine biosynthetic pathway (HBP).^23^ Subsequently, UDP-GlcNAc, along with other nucleotide sugars, is used as glycosyl donors in the process of glycosylation that covalently attach carbohydrate moieties to biomolecules. In the cytosol and nucleus, UDP-GlcNAc is used by O-GlcNAc transferase (OGT) to add O-GlcNAc modifications to proteins, resulting in reversible and regulated cell signaling.^24^ In the endoplasmic reticulum (ER) and Golgi apparatus, UDP-GlcNAc and other nucleotide sugars are used by glycosyltransferases to form complex glycosylation products, which may be further displayed on the cell surface or secreted to the extracellular space.^25^

Because of the importance of UDP-GlcNAc in metabolism and signaling, we have devoted efforts to developing fluorescent UDP-GlcNAc biosensors. We recently reported a green fluorescent biosensor, namely UGAcS, which is an insertion of a circularly permuted green FP (cpGFP) into an inactive mutant of an E. coli UDP-GlcNAc transferase.^26^ UGAcS has the right affinity for monitoring the levels of UDP-GlcNAc in the cytosol. In this manuscript, we describe a parallel effort in our lab, which explores a pBoF-modified circularly permuted red FP (cpRFP) for the specific detection of UDP-GlcNAc (Fig. 2a). The work has resulted in a novel genetically encoded red fluorescent boronolectin, denoted as bapaUGAc, with a suitable affinity for detecting the levels of UDP-GlcNAc in the ER and Golgi.

**Figure 2.**
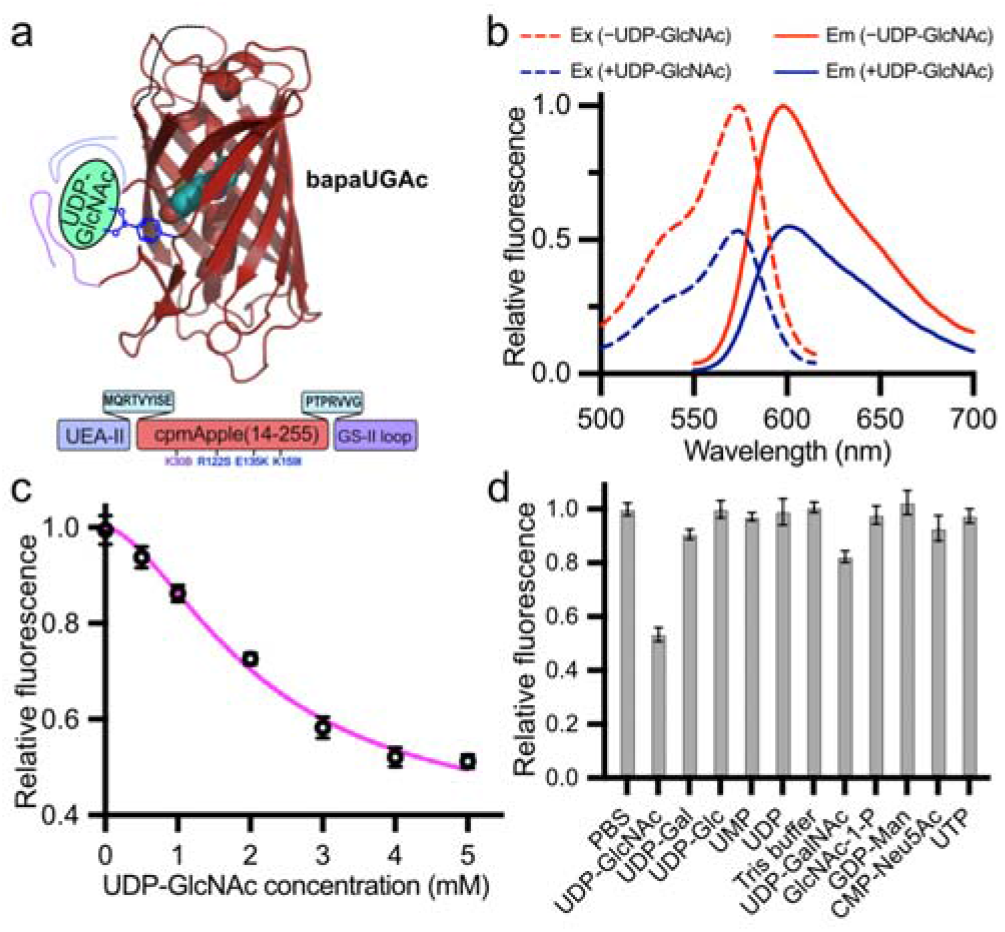
(a) Schematic illustration of bapaUGAc and its interaction with UDP-GlcNAc. The cpRFP chromophore is displayed as cyan spheres. The bottom shows the domain arrangement with the linkers and mutations highlighted, and B represents the pBoF residue. (b) Fluorescence excitation (dashed line) and emission (solid line) spectra of bapaUGAc in the presence (blue) and absence (red) of UDP-GlcNAc. (c) Dose-dependent fluorescence response. (d) Specificity characterization with 1 µM bapaUGAc protein and 5 mM of each compound. Data are presented as mean and s.d. of three technical replicates.

Our biosensor engineering journey started with cpmApple, a cpRFP mutant derived from a previously reported calcium indicator, R-GECO1 (Supporting Information, Fig. S1).^27,28^ Since cpmApple showed relatively low folding efficiency, we randomized the gene using error-prone polymerase chain reactions (EP-PCRs) and screened the library for increased fluorescence in E. coli colonies at 37 °C. We repeated this process for three rounds, resulting in an enhanced cpmApple mutant (ecpApple) with three mutations from cpmApple (Fig. S2). Next, we sought to introduce pBoF into ecpApple. Because we aimed at using carbohydrate binding to modulate fluorescence, we selected two residues (S14 and K30, Figs. S2 & S3) in spatial proximity to the chromophore of ecpApple and site-specifically introduced pBoF to each of the two residues. Among the two mutants, ecpApple with pBoF at residue 30 (ecpApple-K30B) showed a reproducible fluorescence turn-off response to 5 mM UDP-GlcNAc, but the response was not specific since several other carbohydrate analogs could also cause fluorescence changes.

To increase the specificity of the response, we appended a 13-amino-acid peptide sequence derived from a natural N-acetylglucosamine (GlcNAc)-binding lectin, UEA-II (see bapaUGAc0.1 in Fig. S1).^29^ Next, we added several randomized residues to the C-terminus, followed by examining the addition of another peptide sequence derived from natural GlcNAc binding lectins. We tested several options, and a 14-amino-acid loop sequence derived from the GS-II lectin^30,31^ led to the best responsiveness and specificity (bapaUGAc0.3 in Fig. S1). Furthermore, we carried out three additional rounds of error-prone PCRs, and in addition, optimized the linker between the UEA-II-derived peptide and the N-terminus of ecpApple. Our effort resulted in the final bapaUGAc mutant (Figs. 2a, S1, and S2).

The fluorescence excitation and emission peaks of bapaUGAc as a purified protein were at 575 and 600 nm, respectively (Fig. 2b). The fluorescence decreased by ∼50% in response to 5 mM UDP-GlcNAc. We further examined the response using several UDP-GlcNAc concentrations, and the apparent dissociate constant (Kd) was determined to be 2.14±0.29 mM (Fig. 2c). The response displayed some cooperative effect, as the Hill coefficient was 1.7. In addition, we examined the specificity of bapaUGAc among common nucleotide sugars and nucleotides (Fig. 2d). UDP-GlcNAc caused robust fluorescence turn-off, while other tested compounds induced no or only small fluorescence changes. As a negative control, we prepared the ecpApple protein and confirmed its unresponsiveness to UDP-GlcNAc (Fig. S4).

We next explored the use of bapaUGAc to image UDP-GlcNAc level changes in mammalian cells. The Kd of bapaUGAc is too high compared to the concentrations of UDP-GlcNAc in the cytosol. On the other hand, nucleotide sugars are more concentrated in the ER and Golgi,^32^ so we tested whether bapaUGAc could sense UDP-GlcNAc in the ER and Golgi. We used signal peptides to genetically express bapaUGAc in the corresponding subcellular compartments of human embryonic kidney (HEK) 293T cells (Fig. 3a,b). We first treated the cells with GlcN, a potent precursor that can boost the biosynthesis of UDP-GlcNAc (Fig. 3c). As expected, the fluorescence of both ER- and Golgi-localized bapaUGAc decreased in response to GlcN (Fig. 3d,e). Next, we treated the cells with 2-deoxy-d-glucose (2-DG), a glucose analog inhibiting hexokinase (HK) and phosphoglucose isomerase (PGI) (Fig. 3c). The addition of 2-DG caused a robust rise of bapaUGAc signals in the ER or Golgi (Fig. 3 f,g). In contrast, cells expressing ecpApple in the ER or Golgi were unresponsive to GlcN and 2-DG (Fig. 3 d,e,f,g). Collectively, the results support that bapaUGAc can detect UDP-GlcNAc level changes in the ER and Golgi in living mammalian cells in response to metabolic disruption and pharmacological inhibition.

**Figure 3.**
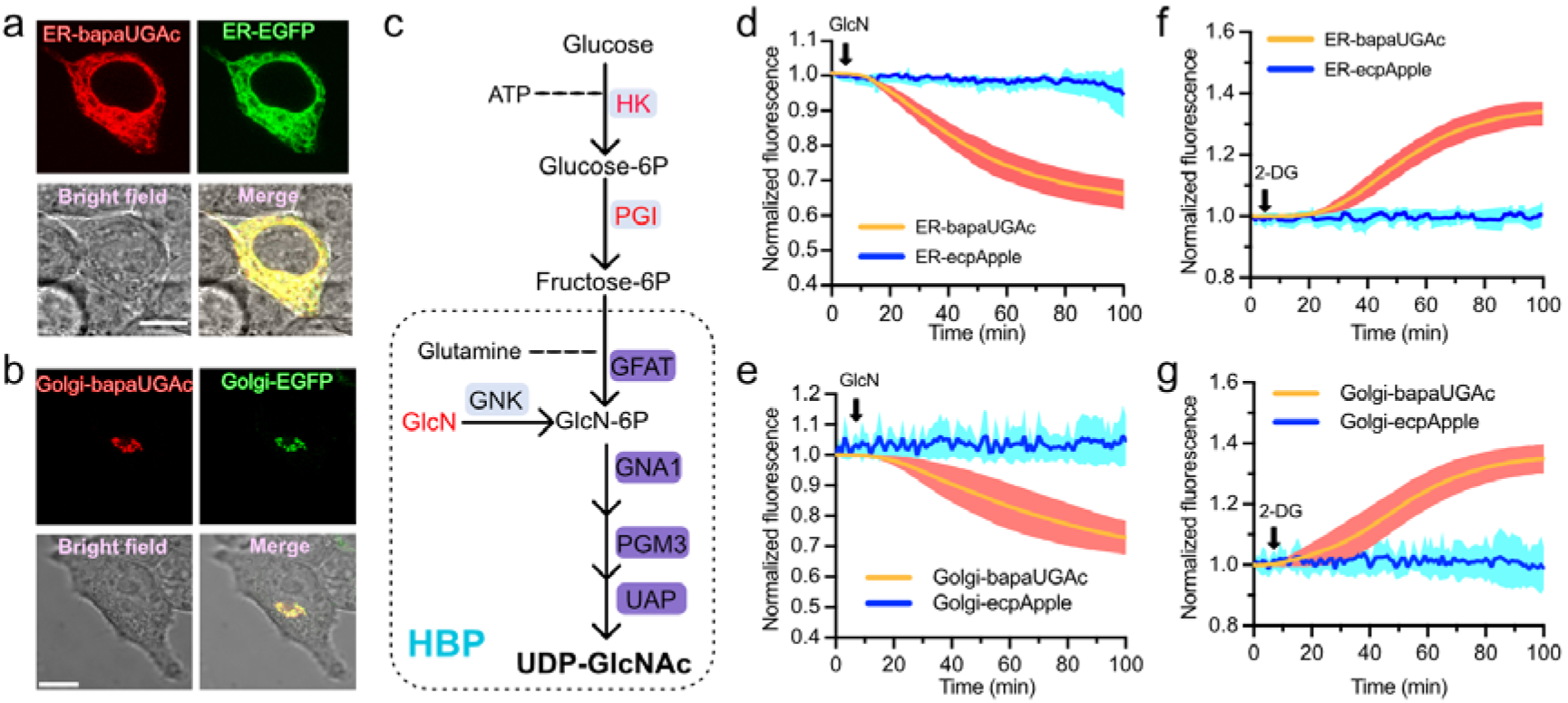
Monitoring UDP-GlcNAc level changes in the ER and Golgi in live mammalian cells using bapaUGAc. (a, b) Co-expression of ER-bapaUGAc and ER-EGFP (a), or Golgi-bapaUGAc and Golgi-EGFP (b) in HEK 293T cells. Scale bar, 10 µm. (c) Schematic illustration of the hexosamine biosynthetic pathway (HBP). Glucosamine (GlcN), as well as hexokinase (HK) and phosphoglucose isomerase (PGI) that are enzymes inhibited by 2-deoxy-d-glucose (2-DG), are highlighted in magenta. (d, e) Time-lapse responses of ER (d) or Golgi (e) localized bapaUGAc in HEK 293T to extracellular addition of GlcN (10 mM) (f, g) Time-lapse responses of ER (f) or Golgi (g) localized bapaUGAc in HEK 293T to extracellular addition of 2-DG (10 mM). Data are presented as mean and s.d. of eight individual cells from three technical replicates.

To visualize UDP-GlcNAc in different subcellular compartments, we paired ER-localized bapaUGAc with our cytosolic UGAcS biosensor. We co-expressed the two biosensors in HEK 293T cells and monitored fluorescence changes in the green and red channels after adding GlcN, a precursor for UDP-GlcNAc synthesis (Fig. 4). Since UGAcS is a fluorescence turn-on sensor, the green fluorescence of UGAcS increased in response to GlcN, indicating an increase in UDP-GlcNAc in the cytosol. Meanwhile, the red fluorescence of bapaUGAc decreased over time, suggesting an accompanied increase in UDP-GlcNAc in the ER. These results support the compatibility of bapaUGAc and UGAcS for dual-color, dual-compartment imaging of UDP-GlcNAc.

**Figure 4.**
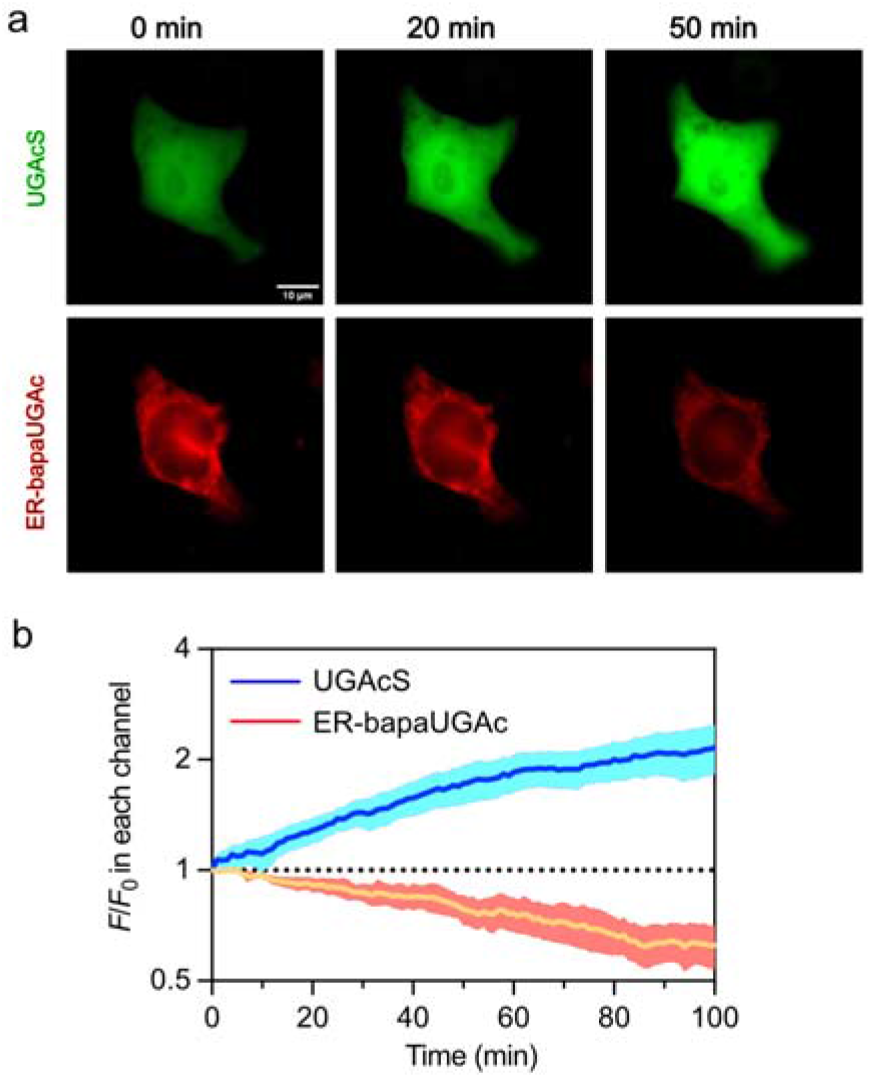
Dual-color imaging of UDP-GlcNAc level changes in the cytosol and ER using green fluorescent UGAcS and red fluorescent bapaUGAc. (a) Representative fluorescence images. Scale bar, 10 µm. (b) Quantitation of the responses of UGAcS and bapaUGAc to extracellular addition of GlcN (10 mM). F/F0 represents the fluorescence intensities of individual cells normalized to the value at starting time. Data are presented as mean and s.d. of 12 individual cells from three technical replicates.

In summary, we have integrated GCE and protein engineering to create a new FP-based boronolectin called bapaUGAc, which can serve as a specific sensor for UDP-GlcNAc. We characterized the functionality of bapaUGAc both in vitro and in live cells. We further achieved dual-color monitoring of UDP-GlcNAc levels in two different subcellular compartments. We expect bapaUGAc to be a valuable research tool for studying UDP-GlcNAc and glycobiology. In addition, our results will catalyze the future development of specific boronolectins for other carbohydrate molecules.

## AUTHOR INFORMATION

### Author Contributions

J.Z., Y.F., and Y.P. engineered the sensors. J.Z. performed characterization experiments with the help of Z.L.. J.Z. analyzed data, prepared figures, and wrote the manuscript. H.-w.A. supervised research, revised figures, and wrote the manuscript.

### Notes

The authors declare no competing financial interest.

## ACKNOWLEDGMENT

Research reported in this publication was supported by the NIH Common Fund Glycoscience Program and the NIH Awards U01CA230817 and R01DK122253. ChatGPT from OpenAI was used to reword and revise some sentences in the manuscript.

