## Supporting Information for "Genetically Encoded Boronolectin as a Specific Red Fluorescent UDP-GlcNAc Biosensor"

### Electronic Supplementary Information

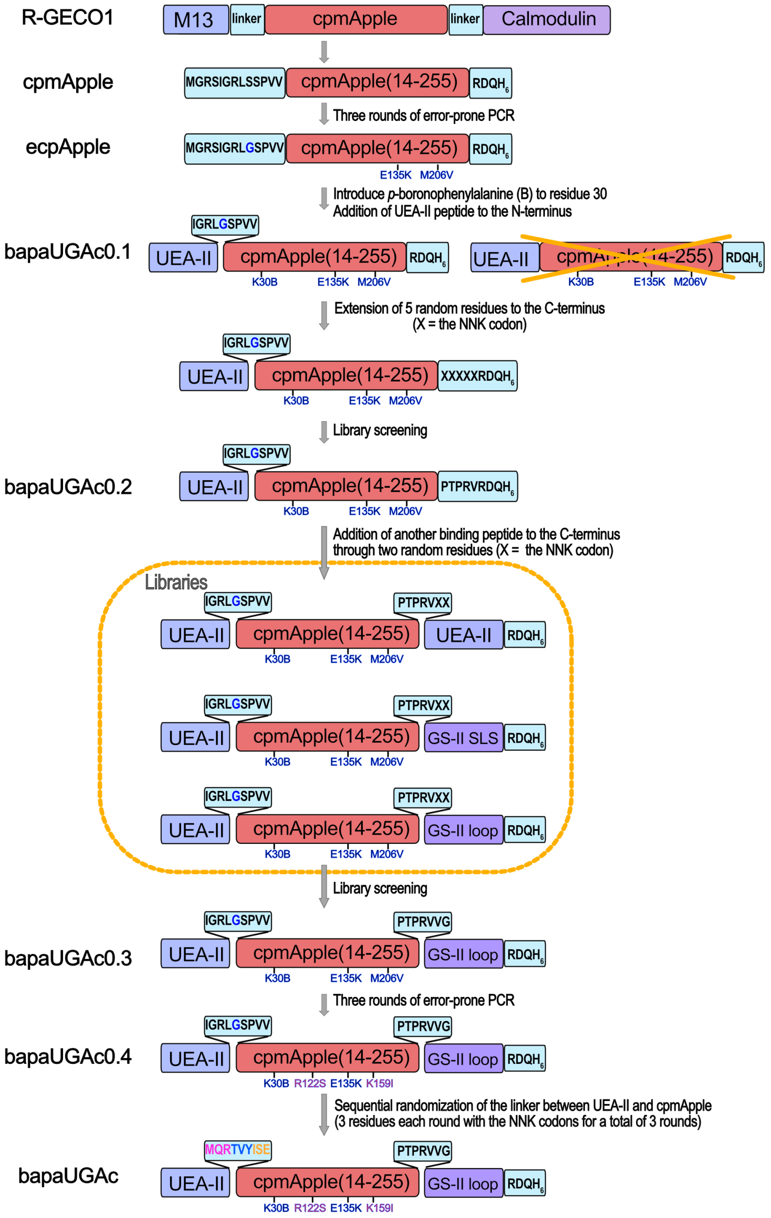


**Figure S1. Illustration of the multistep process to engineer bapaUGAc.** A calcium indicator, R-GECO1, was used to derive a circularly permuted RFP, cpmApple, which further underwent three rounds of error-prone PCRs to derive ecpApple (enhanced cpmApple). Next, *p*BoF was introduced, and a UEA-II-derived GlcNAc-binding peptide was added to the N-terminus. Further, the C-terminal sequences were optimized via screening a 5-residue random library, followed by comparing the addition of the second GlcNAc-binding peptide through two randomized residues. The resultant bapaUGAc0.3 were enhanced via three additional rounds of error-prone PCRs. Finally, the linker between the UEA-II-derived peptide and the N-terminus of ecpApple was optimized to derive bapaUGAc.

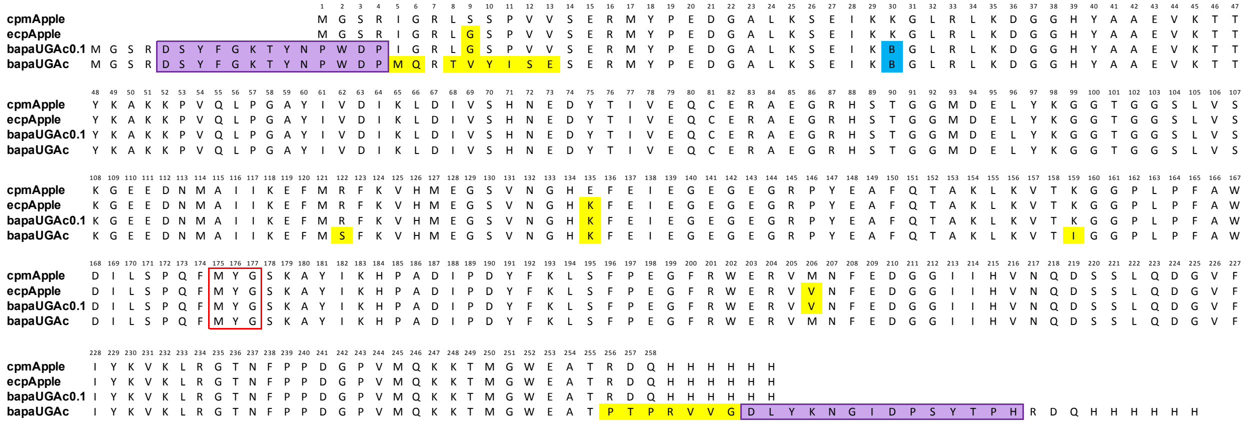

**Figure S2. Alignment of protein sequences of cpmApple, ecpApple, bapaUGAc0.1 and bapaUGAc.** The *p*BoF (B) residue is shaded in blue, and other mutations are shaded in yellow. The RFP chromophore is boxed in red. The UEA-II-derived and GS-II-loop-based GlcNAc-binding peptides are boxed and shaded in magenta.

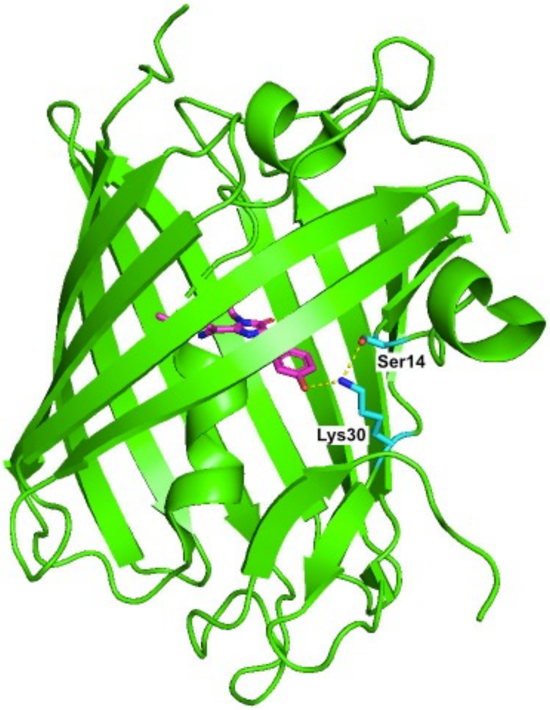


**Figure S3. Illustration of K30 and S14 in cpmApple in relation to the chromophore.** The chromophore is colored in magenta, and K30 and S14 are colored in cyan. H-bonds between them are presented as yellow dashes. The graph is generated based on Protein Data Bank (PDB) entry 4I2Y.[1]

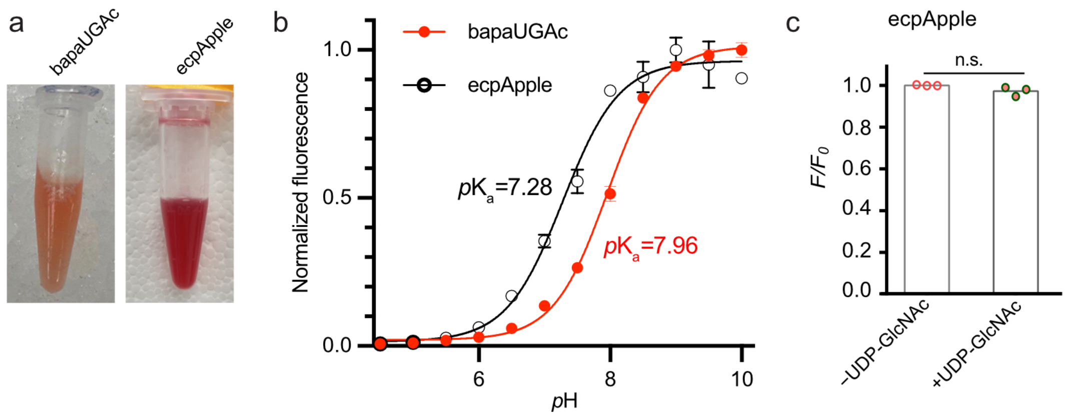


**Figure S4. Additional characterization of bapaUGAc and ecpApple as purified proteins.** (a) Photos of the two purified proteins. (b) *p*H-dependence of the fluorescence of bapaUGAc and ecpApple, showing the apparent $pK_a$ of 7.96 and 7.28, respectively. (c) Fluorescence of ecpApple in the presence and absence of UDP-GlcNAc at *p*H 7.4, supporting that ecpApple can be used as a negative control to verify bapaUGAc responses.

**Table S1. Sequences of oligos used in this study.**

| Name | Sequence (5′→3′) |
|---|---|
| pBAD-modified-cpmApple-F | GGGCTCGAGAATAGGTCGGCTGAGCTCA |
| pBAD-modified-cpmApple-R | GCCAAGCTTAATGATGGTGGTGATGGTGTTGGTCACGCGTAGCC |
| pBAD-F | ATCGCAACTCTCTACTGTTTCTC |
| pBAD-R | CTACTGCCGCCAGGCAAATT |
| Ser14TAG-F | GGCTCACCCGTAGTTTAGGAGCGGATGTACCCC |
| Ser14TAG-R | GGGGTACATCCGCTCCTAAACTACGGGTGAGCC |
| Lys30TAG-F | AAGAGCGAGATCAAGTAGGGGCTGAGGCTGAAG |
| Lys30TAG-R | CTTCAGCCTCAGCCCCTACTTGATCTCGCTCTT |
| pBAD-UEA-II-F | GGCTCGAGAGATTCTTATTTTGGAAAAACTTATAATCCATGGGATCCTATAGGTCGGCTGGGC |
| 5NNK-R | CCAAGCTTAATGATGGTGGTGATGGTGTTGGTCACGMNNMNNMNNMNNMNNCGTAGCCTCCCAGCC |
| cpmApple-UEA-II-F | ACGCCTACGCCTCGGGTGNNKNNKGATTCTTATTTTGGAA |
| cpmApple-UEA-II-R | TTCCAAAATAAGAATCMNNMNNCACCCGAGGCGTAGGCGT |
| UEA-II-R | GCCAAGCTTAATGATGGTGGTGATGGTGTTGGTCACGAGGATC |
| pBAD-UEA-II-3NNKs-F | GGCTCGAGAGATTCTTATTTTGGAAAAACTTATAATCCATGGGATCCTNNKNNKNNKCTGGGCTCACCCGTA |
| MAH-ER-F | TCCCAGGTCCAACTGCACGGAAGCTTGCCACCATGCTGCTATCCGTGCCGCTG |
| ER-bapaUGAc-F | TCGGCCTGGCCGCAGCTGACGAATTCGATTCTTATTTTGGAAAAAC |
| ER-bapaUGAc-R | GTTTTTCCAAAATAAGAATCGAATTCGTCAGCTGCGGCCAGGCCGA |
| MAH-bapaUGAc-KDEL-R | GCTGATCAGCGGGTTTAAACGGGCCCCTAGAGTTCGTCCTTTTGGTCACGATGAGGAGT |
| MAH-Golgi-F | TCCCAGGTCCAACTGCACGGAAGCTTGCCACCATGAGGCTTCGGGAGCCGCTC |
| Golgi-bapaUGAc-R | TTTTCCAAAATAAGAATCGAATTCGGCCCCTCCGGTCCGG |
| Golgi-bapaUGAc-F | CCGGACCGGAGGGGCCGAATTCGATTCTTATTTTGGAAAA |
| MAH-bapaUGAc-R | GCTGATCAGCGGGTTTAAACGGGCCCCTATTGGTCACGATGAGGAGT |

### EXPERIMENTAL METHODS

#### Key reagents and methods

This study used the racemic form of the *p*-borono-DL-phenylalanine (*p*BoF) amino acid purchased from Syntonix (Syntonix, Waltham, USA). Uridine diphosphate *N*-acetylglucosamine (UDP-GlcNAc), uridine diphosphate *N*-acetylgalactosamine (UDP-GalNAc), uridine diphosphate Galactose (UDP-Gal), cytidine monophosphate sialic acid (CMP-Sia), GDP-fructose, GDP-mannose, UTP, and D-glucosamine hydrochloride (GlcN) were purchased from MilliporeSigma (St. Louis, MO, USA). UDP, 2-deoxy-D-glucose (2-DG), and L-arabinose were purchased from Cayman Chemical (Ann Arbor, Michigan, USA). Synthetic DNA oligos were ordered from Integrated DNA Technologies (Coralville, Iowa, USA). DNA sequences were confirmed with Sanger sequencing performed by Eurofins Genomics (Louisville, KY, USA). pEvol-*p*BoF was a gift from Dr. Peter Schultz (Scripps Research).[2] pMAH-POLY and pcDNA-UGAcS were from our previous studies.[3, 4]

#### Engineering of bapaUGAc.

The cpmApple gene fragment was amplified from R-GECO1 using oligos pBAD-modified-cpmApple-F and pBAD-modified-cpmApple-R (see **Table S1** for oligo sequences).[5, 6] The PCR product was digested with Xho I and Hind III and inserted into a pre-digested, modified pBAD plasmid, resulting in pBAD-cpmApple (**Fig. S2**).

To further improve the brightness of cpmApple, error-prone PCRs (EP-PCRs) were performed using oligos pBAD-F and pBAD-R, according to a previously described procedure.[7] Next, the PCR product was digested with Xho I and Hind III and inserted into the abovementioned, modified pBAD plasmid. The DNA library was used to transform *E. coli* DH10B electrocompetent cells. Cells were plated on 2×YT agar plates supplemented with 100 μg/mL ampicillin and 0.02% (w/v) L-arabinose. Next day, colonies on the agar plates were imaged using a customized imaging system equipped with a Dolan-Jenner Mi-LED Fiber Optic light source, appropriate excitation and emission filters in Thorlabs motorized filter wheels, and a QSI 628 CCD camera. Colonies with higher fluorescence were selected and cultured in 1 mL 2×YT supplemented with 100 μg/mL ampicillin in the wells of a 96-well deep-well plate shaken at 250 r.p.m. and 37 °C overnight. Next, 1 mL of fresh 2xYT media supplemented with 100 μg/mL ampicillin and 0.04% (w/v) L-arabinose was added to the overnight starter culture, and the new mixtures were incubated for additional 48 hr at 250 r.p.m. and 30 °C. Centrifugation at 3800 ×g was used to pellet cells, which were further lysed with 300 μL of a bacterial lysis buffer (5 mg/mL octyl glucoside, 0.1 mg/mL chicken egg lysozyme, and 0.2 U/mL Benzonase in Tris-HCl, pH 8) by shaking at 200 r.p.m. on ice for 1 hr. The fluorescence of the cell lysates was measured on a BioTek Synergy Mx microplate reader with the excitation wavelength at 575 nm and the emission wavelength at 600 nm. Plasmid DNAs were extracted from colonies showing high fluorescence and sequences were determined via Sanger sequencing. Three rounds of directed evolution were carried out to derive ecpApple.

Next, site-directed mutagenesis was used to introduce the amber (TAG) codon to residues 14 and 30 of ecpApple, respectively. Briefly, oligos pBAD-F and Ser14TAG-R, or Ser14TAG-F and pBAD-R, were used to amplify fragments of ecpApple. These fragments were assembled with pBAD-F and pBAD-R in an overlap PCR. The PCR product was digested with Xho I and Hind III and inserted into the pBAD plasmid, resulting in pBAD-ecpApple-S14TAG. Similarly, oligos pBAD-F, Lys30TAG-R, Lys30TAG-F and pBAD-R were used to generate pBAD-ecpApple-K30TAG. The resultant pBAD plasmids, along with pEvol-*p*BoF,[2] were used to co-transform DH10B electrocompetent cells. Protein purification and initial substrate specificity test were carried out, according to the procedures presented in the next section.

pBAD-ecpApple-K30TAG was chosen for further engineering. To derive bapaUGAc0.1, oligos pBAD-UEA-II-F and pBAD-R were used to amplify the ecpApple-K30TAG gene fragment and add an N-terminal 13-amino-acid peptide sequence (DSYFGKTYNPWDP) derived from the *Ulex europaeus* agglutinin II (UEA-II) lectin. The PCR product was inserted into pBAD as described above. Next, oligos pBAD-F and 5NNK-R were used to amplify bapaUGAc0.1 and add five randomized residues to the C-terminus. The library was screened using a procedure similar to the engineering of ecpApple, except that both high brightness and UDP-GlcNAc responsiveness were used as the selection criteria. This procedure led to bapaUGAc0.2.

Furthermore, three additional peptide sequences derived from natural GlcNAc-binding lectins were added to the C-terminus of bapaUGAc0.2 via two additional randomized residues. Briefly, to add the second copy of the 13-amino-acid peptide sequence from UEA-II, oligos pBAD-F and cpmApple-UEA-II-R were used to amplify a gene fragment from pBAD-bapaUGAc0.2, and oligos cpmApple-UEA-II-F and UEA-II-R were used in a PCR without additional templates to generate a short fragment; next, the two fragments were assembled using oligos pBAD-F and UEA-II-R via an overlap PCR. The assembled fragment was digested with Xho I and Hind III and inserted into the pBAD plasmid. Similar procedures were used to add peptide sequences derived from the *Griffonia simplicifolia* GS-II lectin, and these sequences were previously reported to interact with GlcNAc.[8, 9] GS-II SLS refers to a 27-amino acid sequence (IVFCEFDLYKNGIDPSYTPHLGINVNQ), while GS-II loop refers to a 14-amino-acid sequence (DLYKNGIDPSYTPH). Screening of these three libraries for high brightness and UDP-GlcNAc responsiveness led to bapaUGAc0.3, which is essentially bapaUGAc0.2 linked to the N-terminus of GS-II loop via a two-amino-acid linker.

From bapaUGAc0.3, three rounds of directed evolution based on EP-PCRs were performed by following the procedure described above. The best-selected clone was bapaUGAc0.4. To further optimize the linker between the N-terminal UEA-II-derived peptide and cpmApple, oligos pBAD-UEA-II-3NNKs-F and pBAD-R were used to amplify bapaUGAc0.4 and randomize the first three residues of this 9-amino-acid linker. The PCR product was digested by Xho I and Hind III and then inserted into the predigested pBAD plasmid. Screening of the library identified an improved mutant, which was used for further engineering. Next, similar procedures were used to randomize the middle and the last three residues of this 9-amino-acid linker. Screening of the libraries led to the final mutant, namely bapaUGAc.

**Protein purification and *in vitro* characterization.**

To express the bapaUGAc protein, pBAD-bapaUGAc and pEvol-*p*BoF were used to co-transform DH10B electrocompetent cells. A single colony was used to inoculate 3.0 mL of 2×YT media supplemented with 100 μg/mL ampicillin and 50 μg/mL chloramphenicol at 220 r.p.m. and 37 °C overnight. The saturated starter was added into 300 mL of Terrific Broth medium (TB) supplemented with the same concentrations of antibiotics mentioned above. When the optical density (OD) at 600 nm reached 1.0, 0.02% (w/v) L-arabinose and 2 mM *p*BOF were added. The culture was incubated at 220 r.p.m. and 30 °C for additional 48 hr. Cells were next pelleted, resuspended in 1× phosphate-buffered saline (PBS, pH 7.4), and lysed on ice by sonication. The lysate was clarified with centrifugation at 16000 ×g and 4 °C for 30 min. The supernatant was subjected to Ni-NTA agarose bead (Pierce, Rockford, IL) affinity purification. Proteins in ~ 4 mL of 300 mM imidazole-containing buffer were next injected into a HiLoad 16/600 Superdex 200 pg size-exclusion column on an AKTA protein purification system (Cytiva). 1× PBS was used for elution, and the eluted proteins were kept at 4 °C. Protein concentration was determined using the alkali denaturation method by assuming the extinction coefficient of the denatured chromophore to be 44,000 $M^{-1}cm^{-1}$.[10]

Fluorescence spectra of the purified protein (1 μM) in the PBS buffer in the presence and absence of 5 mM UDP-GlcNAc were measured using a BioTek Synergy Mx Microplate Reader. To record the excitation spectra, the emission wavelength was set at 640 nm while the excitation was scanned from 500 nm to 620 nm. To record the emission spectra, the excitation wavelength was set at 530 nm while the emission was scanned from 550 to 700 nm. Next, to determine the dose-dependent response of bapaUGAc to UDP-GlcNAc, 1 μM protein was incubated with PBS or UDP-GlcNAc in PBS at final concentrations of 0.5, 1, 2, 3, 4, and 5 mM, respectively. In addition, to examine the indicator's specificity, the freshly prepared protein (1 μM) was incubated with chosen chemicals in PBS at a final concentration of 5 mM. Endpoint fluorescence intensity measurement was carried out 5 min after mixing all reagents with excitation and emission set at 570 and 600 nm, respectively.

**Construction of mammalian expression plasmids.**
To generate a plasmid for bapaUGAc expression in the ER, oligos ER-bapaUGAc-F and bapaUGAc-KDEL-R were used to amplify the bapaUGAc gene from pBAD-bapaUGAc. This step also added a C-terminal ER retention sequence (KDEL). Meanwhile, pMAH-ER-F and ER-bapaUGAc-R were used in a PCR without additional templates to generate a short fragment encoding the N-terminal ER signal peptide (MLLSVPLLLGLLGLAAAD) derived from calreticulin. The two resultant DNA fragments, along with a pMAH plasmid predigested with Hind III and Apa I, were subjected to a three-fragment Gibson assembly reaction, resulting in the pMAH-ER-bapaUGAc plasmid. A similar procedure was used to construct a plasmid for bapaUGAc expression in the Golgi apparatus. Briefly, oligos Golgi-bapaUGAc-F and MAH-bapaUGAc-R were used to amplify the bapaUGAc gene from pBAD-bapaUGAc. Oligos MAH-Golgi-F and MAH-Golgi-bapaUGAc-R were used to amplify an 81-amino-acid Golgi signal peptide derived from human β-1,4-galactosyltransferase 1, and HeLa cell cDNAs were used as the PCR template. The resultant fragments, along with pMAH predigested with Hind III and Apa I, were again assembled via Gibson assembly, resulting in the pMAH-Golgi-bapaUGAc plasmid. Negative control plasmids, pMAH-ER-ecpApple and pMAH-Golgi-ecpApple were generated using similar procedures from pBAD-ecpApple.

**Mammalian cell culture, transfection, and live cell imaging.**
Human Embryonic Kidney (HEK) 293T cells were cultured in Dulbecco's Modified Eagle's Medium (DMEM) containing 4.5 g/L glucose supplemented with 10% fetal bovine serum (FBS) in the humidified incubator containing 5% $CO_2$ at 37 °C. HEK 293T cells were seeded in 35-mm tissue culture dishes with glass coverslips at the bottom. Transfection was performed on the second day when cell confluency reached ~30%. Briefly, 1.5 μg pMAH-POLY and 1.5 μg pMAH-ER-bapaUGAc (or pMAH-Golgi-bapaUGAc) were added to 7.5 μg of PEI (polyethyleneimine, linear, M.W. 25 kDa) in 600 μL Opti-MEM (Gibco). For the dual-color imaging experiment, 1.5 μg pMAH-POLY, 1.5 μg pMAH-ER-bapaUGAc, 1 μg pcDNA-UGAcS, and 9 μg of PEI were used. The plasmid-PEI mixture was incubated at room

temperature for 30 min before being added to HEK 293T cells in culture dishes. 16 hr later, fresh DMEM media containing 10% FBS and 2 mM *p*BoF were used to replace the transfection media. After another 24 hr, to deplete intracellular free *p*BoF amino acid, the culture media were again replaced with fresh DMEM containing 10% FBS but no *p*BoF. Imaging was performed 12 hr later. Cells were rinsed with a mammalian cell imaging buffer (140 mM NaCl, 5 mM KCl, 1 mM $CaCl_2$, 0.4 mM $MgSO_4$, 0.5 mM $MgCl_2$, 0.3 mM $Na_2HPO_4$, and 0.4 mM $KH_2PO_4$, 6 mM D-glucose, and 20 mM HEPES, pH 7.4) three times and kept in this buffer at room temperature for 30 min for stabilization of metabolism. Cells were next imaged using a Leica DMi8 inverted microscope equipped with a Leica EL6000 light source, a TRITC filter cube (545/25 nm bandpass excitation and 605/70-nm bandpass emission), and a Photometrics Prime 95B sCMOS camera. To perturb the intracellular UDP-GlcNAc level, cells were treated with either 10 mM GlcN or 10 mM 2-DG pre-dissolved in the imaging buffer. Images were analyzed using the ImageJ software. Backgrounds were subtracted using the software by setting the rolling ball radius to 100 pixels. Intensity means of randomly selected cells were used for analysis. The fluorescence changes were calculated by normalizing fluorescence intensities (F) to the value at starting time ($F_0$), and the $F/F_0$ ratios were plotted against time.
